# Cell wall thickness has phylogenetically consistent effects on photosynthetic nitrogen-use efficiency of terrestrial plants: a meta-analysis

**DOI:** 10.1101/2022.08.23.505027

**Authors:** Dan-dan Liu, Tiina Tosens, Dong-liang Xiong, Marc Carriquí, You-cai Xiong, Wei Xue

## Abstract

Leaf photosynthetic nitrogen-use efficiency (PNUE) diversified significantly among C_3_ species. However, morpho-physiological mechanisms and interrelationships forming PNUE remain unclear on the evolutionary time scale. In this study, we compiled a novel extensive matrix of morpho-anatomical and physiological traits of leaf in 679 C_3_ species ranging from bryophytes to angiosperms to understand the intricacy of interrelationships underlying the variations in PNUE. We found that LMA, mesophyll cell wall thickness (T_cwm_), Rubisco N allocation fraction (P_R_), and mesophyll conductance (g_m_) together interpreted 83% of variations in PNUE, with P_R_ and g_m_ accounting for 65% of those variations. However, the P_R_ effects were species-dependent on g_m_; that is, the contribution of P_R_ on PNUE was extensively significant in high-g_m_ species in comparison to low-g_m_ species. Standard major analysis (SMA) and path analysis suggested a weak correlation between PNUE and LMA, whereas the SMA correlation for PNUE–T_cwm_ was strong. The P_R_ was inversely proportional to T_cwm_, which was similar to the relationship between g_m_ and T_cwm_ (p-value < 0.01), so that the internal CO_2_ drawdown from intercellular airspace to carboxylaton sites was relatively conservative over a wide range of T_cwm_. Collectively, the coordination of changes in P_R_ and g_m_ connecting T_cwm_ suggested the complex physiological mechanisms mediated by T_cwm_ modulating PNUE across contrasting plant groups.

## Introduction

Trait-based ecology is an emerging discipline, aiming to predict functional diversity of population, communities, and ecosystems using the generalizable, functional properties of individual organisms (De Bello *et al*., 2021). Leaf photosynthetic nitrogen-use efficiency (PNUE), defined as the ratio of light-saturated photosynthesis to leaf nitrogen content, is a fundamental process of the leaf economics spectrum (LES) (Wright *et al*., 2005). Significant variations were observed in the PNUE economic trait. In spermatophyte, there is a correlation between PNUE and leaf mass per area (LMA); species with high LMA almost invariably have low PNUE (Reich *et al*., 1998; Wright *et al*., 2005; Onoda *et al*., 2017). The LMA, as a complex leaf morphologic trait, could not explain inter-specific PNUE variations (Shipley *et al*., 2006; Warren and Adams, 2006; Hidaka and Kitayama, 2009; Onoda *et al*., 2017; Flexas and Carriquí, 2020; Luo *et al*., 2021). The morpho-physiological mechanisms and interrelationships underlying the PNUE are attracting ever-increasing attention owning to the ecological significance of the inter-specific variations of PNUE in relation to relative growth rate and crop yield or for the sake of realistic ecosystem modeling (Poorter *et al*., 1990; Garnier *et al*., 1995; Reich *et al*., 1998; Oguchi *et al*., 2016; Yin *et al*., 2019).

Previous studies demonstrated that variations in PNUE resulted from interactive mechanisms of several physiological and anatomical traits of foliage (Hikosaka, 2004, 2010; Wright *et al*., 2005; Onoda *et al*., 2017; Flexas and Carriquí, 2020). Specifically, nitrogen allocation into photosynthetic apparatus and mesophyll conductance for internal CO_2_ diffusion (g_m_) were important. In spermatophyte, experimental studies observed an inverse relationship between LMA and the fraction of nitrogen allocated to photosynthetic proteins. With increasing LMA in ten dicotyledonous species, the photosynthetic N allocation fraction decreased from 50% to 38%, with a significant decrease in the Rubisco N allocation fraction (P_R_) (Poorter and Evans, 1998). Hikosaka *et al*. (1998) analyzed an annual herb and an evergreen tree, reporting a higher P_R_ of 23.2% in the low-LMA herb in comparison to the high-LMA tree. Similar results were also observed in other studies where the high-LMA species and genotypes exhibited a reduced P_R_ (Warren and Adams, 2000; Ripullone *et al*., 2003; Onoda *et al*., 2004; Pons and Westbeek, 2004; Takashima *et al*., 2004; Onoda *et al*., 2017; Gvozdevaite *et al*., 2018; Lei *et al*., 2021a). A meta-analysis in spermatophyte performed by Onoda *et al*. (2017) attributed the negative relationship between PNUE and LMA to the trade-off mechanisms between photosynthesis and structural persistence (hereinafter referred to LMA-mediated function-structure tradeoff in spermatophyte), which means that species at the high-end of the LMA axis were more likely to preserve a greater fraction of leaf nitrogen in structural tissues, resulting in lower P_R_ and PNUE. The PNUE variation from the low-end LMA axis to the high-end was associated with increased N investments in structural tissue. Recently, a study that quantified the negative P_R_*–*LMA relationship in spermatophyta has supported it (Luo *et al*., 2021).

An inevitable result of the construction cost for structural tissues is a decrease in g_m_ (Onoda *et al*., 2017; Flexas and Carriquí, 2020), with a tendency for g_m_ to decline from the low- to the high-end of the LMA axis in spermatophyta. In ferns and mosses, the observed variations in PNUE were strongly associated with g_m_ (Tosens *et al*., 2016; Carriquí *et al*., 2019). Ferns and mosses with similar LMA as seed plants clustering at the low-end of the LMA axis exhibited substantially lower g_m_ and PNUE (Carriquí *et al*., 2015; Tosens *et al*., 2016; Zhu *et al*., 2016; Z. Wang *et al*., 2017), which indeed was in contradiction to the negative PNUE–LMA pattern on which the LMA-based function-structure tradeoffs were dependent. In contrast to the critical roles of g_m_ in PNUE in mosses and ferns, several studies in seed plants discovered that g_m_ did not contribute to PNUE variations in 11 evergreen tropical and temperate species that differed in LMA (Bahar *et al*., 2018) as well as in 26 wild and 31 domesticated cotton genotypes with significant variations in LMA (Lei *et al*., 2021b). It suggested that the LMA-based function-structure tradeoff paradigm developed by Onoda *et al*. (2017) might have a limited capability of addressing the underpinnings of PNUE variations across the phylogeny of land plant.

Bryophytes and pteridophytes are significant from an evolutionary perspective owning to their pioneering positions in the land transition of photosynthetic life; they are currently attracting considerable attention among researchers owning to their great potential in biotechnology and bioindication (Preston and Hill, 1999; Popper and Fry, 2003; Wang *et al*., 2013; Gago *et al*., 2019; Flexas and Carriquí, 2020). Concerning the above mentioned disputes, earlier studies attributed diversions of paired-trait relationships among major plant groups to leaf habits or height-based economics spectrum (Zhao *et al*., 2017; Li *et al*., 2022) or microhabitat acclimations (Keenan and Niinemets, 2016; Anderegg *et al*., 2018) or unknown fundamental difference (Waite and Sack, 2009; Z. Wang *et al*., 2017). Until now, LMA has not been well understood in terms of its role in PNUE variations, its correlations with g_m_ and P_R,_ as well as the mechanisms involved in g_m_ and P_R_ functioning across major plant groups and phylogenetic interrelationships.

The g_m_ is a highly complex trait that mediates liquid phase diffusion and is highly sensitive to leaf anatomy and biochemical traits (e.g., effective porosity of the cell wall-*p*_i_, the CO_2_ permeability of membranes, carbonic anhydrase activity, etc.) (Evans *et al*., 2009; Kaldenhoff, 2012; Tosens *et al*., 2012; Han *et al*., 2018; Gago *et al*., 2019; Carriquí *et al*., 2020; Evans, 2021; Flexas *et al*., 2021), which consolidate attempts to scale PNUE with g_m_ (Warren and Adams, 2006). Pleiotropic compensatory changes (e.g., one advantageous factor compensates for another drawback) might exert distinct roles when comparing PNUE across a few species or species occupying the same habitat in which resource use strategies tend to converge (Evans *et al*., 2009; Anderegg *et al*., 2018; Lei *et al*., 2021b; Nadal *et al*., 2021). In this study, a global extensive database was compiled, including leaf morphological, anatomical, and physiological traits of terrestrial C_3_ species ranging from bryophytes to angiosperm. Data in C_4_ species that have distinct photosynthetic mechanisms than C_3_ species were not the major concern in this study. Data analysis and presentation in *Results* section were performed within major taxonomic group (i.e., bryophytes, pteridophytes, gymnosperm, and angiosperm) and across taxonomic groups, helping clarify the scales at which our results were applicable. Specifically, we aimed to address three questions as follows:

1. What is the evolutionary pattern of PNUE from bryophytes to angiosperms? Are PNUE variations primarily associated with LMA across major taxonomic groups?
2. What are the crucial morpho-anatomical traits in shaping PNUE variations across major taxonomic groups?
3. What are the interrelationships between these important morpho-anatomical traits and PNUE across major taxonomic groups?

## Results

### The evolutionary pattern of PNUE and its relationship with LMA

Significant phylogenetic signals were observed for the LMA, PNUE, A_mass_, N_m_, V_cmax_, J_max_, P_R_, g_m_, DeltaC, and T_cwm_ (Table S1). Initially, we mapped the LMA phylogenetic pattern and its relationship with PNUE, A_mass_, and N_m_ as LMA was considered an important factor influencing the PNUE economic trait in seed plants (Hikosaka, 2004; Wright *et al*., 2005; Onoda *et al*., 2017). The mean of LMA cluster data largely increased from bryophytes to pteridophytes (TukeyHSD, p-value = 0.07) (Figure 1a). No significant differences were observed in mean LMA between pteridophytes and angiosperms (p-value = 0.27, Figure 1a). The mean LMA in gymnosperms was significantly higher in comparison to the other three phyla (p-value < 0.01). The mean PNUE values from the ANOVA revealed significant differences across the four phyla (Figure 1b), with the angiosperms having the highest PNUE values (p-value < 0.05) and the other three phyla having similar PNUE values (p-value > 0.1). Significantly, PNUE was lower in bryophytes at the same LMA as seed plants and marginally lower in pteridophytes (Figure 1c). The LES SMA regression (fit SMA model to all data in the four phyla) exhibited a low r^2^ at 0.1 owning to a significant deviation of bryophytes data from the LES SMA regression line (Figure 1c, SMA ANOVA in intercept: p-value < 0.01). Improved performance in the LES SMA regression for A_mass_–LMA was evident (r^2^ = 0.33); whereas all A_mass_ values in bryophytes significantly deviated from the LES SMA regression line primarily owning to major differences in intercept (Figure 1d, SMA ANOVA in intercept: p-value < 0.01). The LES SMA regression line and the phylum SMA regression lines (fit SMA model to data of each phylum) for the N_m_–LMA relationship were in close proximity to one another (Figure 1e), demonstrating significantly higher r^2^ = 0.53. Inequality in the number of sampled species among the four phyla did not distort the SMA correlation results. It was suggested that variations in PNUE through plant phylogeny were weakly associated with LMA primarily owning to large discrepancies in slope and intercept of the A_mass_–LMA relationship among major taxonomic groups.

**Figure 1.**
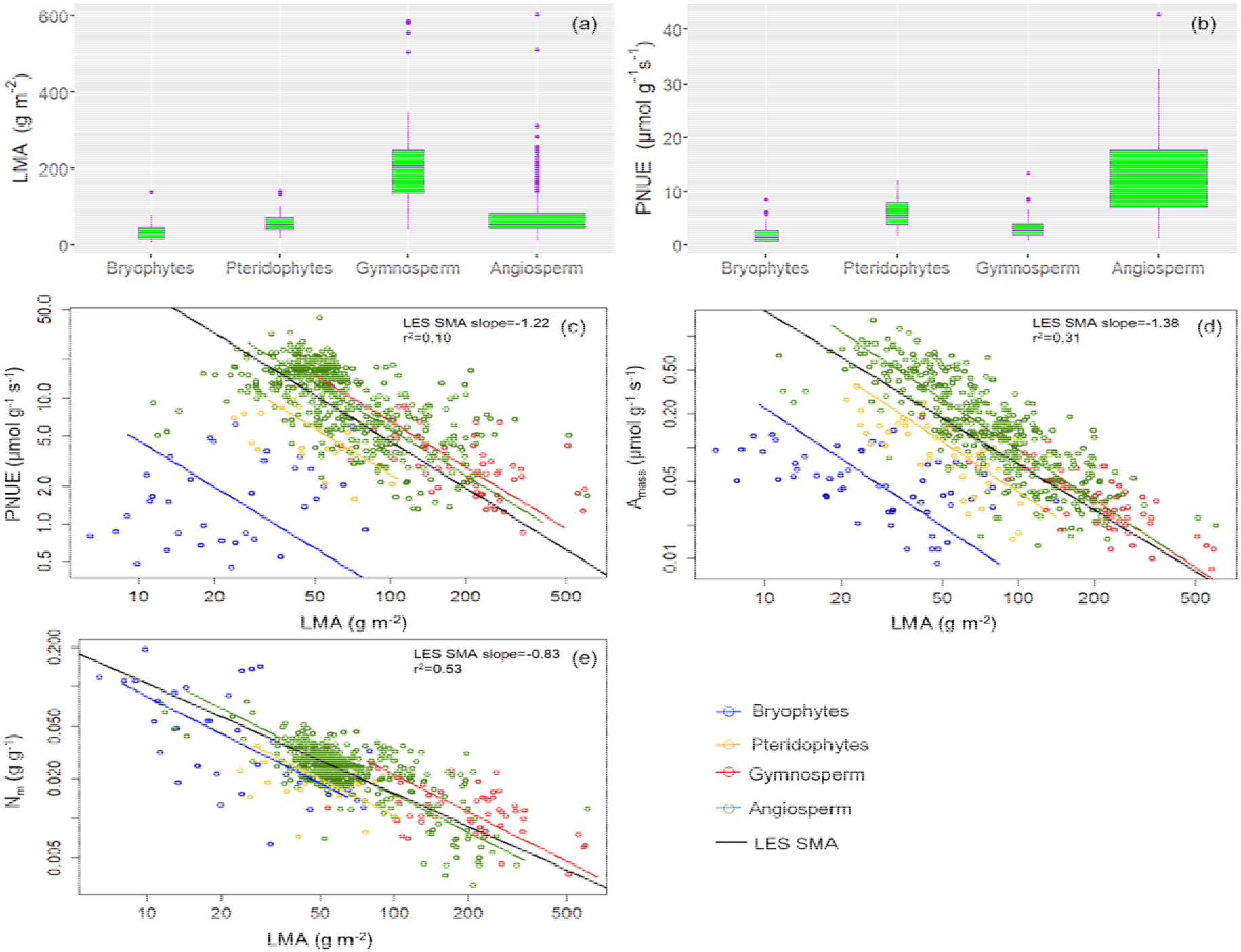
The phylogenetic patterns of leaf mass per area (LMA), (a) and photosynthetic N-use efficiency (PNUE), (b) as well as the paired-trait relationships between PNUE, mass-based photosynthesis capacity (A_mass_), leaf nitrogen content per leaf mass (N_m_), and LMA. (c) PNUE–LMA, (d) A_mass_–LMA, and (e) N_m_–LMA. The black line represents the regression of standard major analysis (SMA) fit to all data in the four phyla (LES SMA), and the other lines represent regressions of SMA fit to all data of each phylum. The r^2^ and slope values were elucidated for the LES SMA regression. The boxes were drawn with widths proportional to the square-roots of the number of observations in each phylum. The same symbol legends for bryophytes, pteridophytes, gymnosperms, and angiosperms as in Figure 1 were used in other figures unless otherwise indicated.

### Effects of photosynthetic nitrogen allocation and g_m_ on photosynthesis and PNUE variations

Although no significant correlation was observed for PNUE–N_m_ (LES SMA r^2^ = 0.002, p-value = 0.32), the LES SMA fit for PNUE–A_mass_ exhibited r^2^ as high as 0.58 (Figure S1), indicating that PNUE changing pattern along the LMA axis was strongly shaped by A_mass_. Therefore, quantitative correlations between important physiological traits and A_mass_ were explored in this section. A positive LES SMA regression line for the A_mass_–V_cmax_ relationship exhibited r^2^ = 0.68, which overlapped the four phylum regression lines efficiently (Figure 2a, SMA ANOVA in slope: p-value = 0.07). A closer correlation of the LES SMA regression fit to A_mass_–g_m_ was evident with r^2^ at 0.74, showing no significant differences against phylum regression lines (Figure 2b, SMA ANOVA in slope: p-value = 0.9). The V_cmax_ was closely related to J_max_ with r^2^ as high as 0.89 (Figure 2c), close to four phylum regression lines (SMA ANOVA in slope: p-value = 0.04). The relationship between g_m_ and V_cmax_ exhibited relatively weaker correlation strength in comparison to V_cmax_–J_max_ (r^2^ = 0.5, Figure 2d).

**Figure 2.**
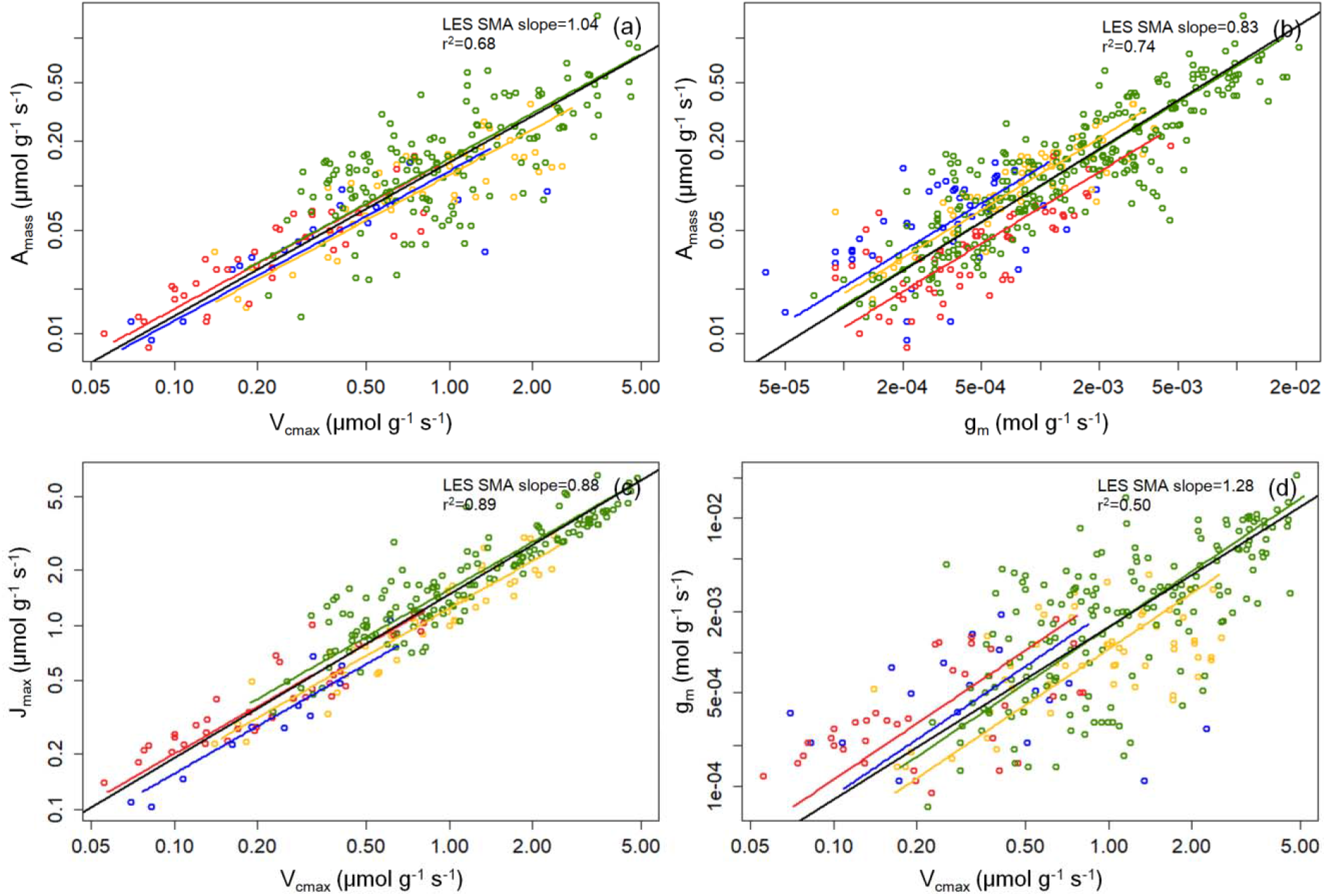
Paired-trait correaltions between the mass-based photosynthesis (A_mass_), the mass-based maximum carboxylation rate (V_cmax_), the mass-based mesophyll conductance (g_m_), and the mass-based maximum electron transport rate (J_max_). (a) A_mass_–V_cmax_, (b) A_mass_–g_m_, (c) V_cmax_–J_max_, and (d) g_m_–V_cmax_. The black line represents the regression of standard major analysis (SMA) fit to all data in the four phyla (LES SMA) and the other lines represents the regressions of SMA fit to all data of each phylum. The r^2^ and slope values were elucidated for the LES SMA regression.

The A_mass_ was significantly regulated by V_cmax_ which was, in turn, related to P_R_ (LES SMA r^2^ = 0.62). According to Onoda *et al*. (2017), DeltaC might be an important factor influencing PNUE variations in seed plants. Hence, PNUE relationships with P_R_, g_m_, and DeltaC were quantified in Figure 3. Across the four phyla, we observed substantial correlation strength between PNUE and P_R_ (r^2^ = 0.45, Figure 3a). The PNUE–g_m_ correlation had a higher r^2^ (0.52, Figure 3c) in comparison to P_R_. According to path analysis, two traits explained approximately 65% of PNUE variations. Across the four phyla, there was no discernible correlation between PNUE and DeltaC (r^2^ = 0.04, Figure 3c). The r^2^ value was considerably lower for the P_R_–g_m_ relationship (r^2^ = 0.20, Figure 3d), indicating a weak correlation between them.

**Figure 3.**
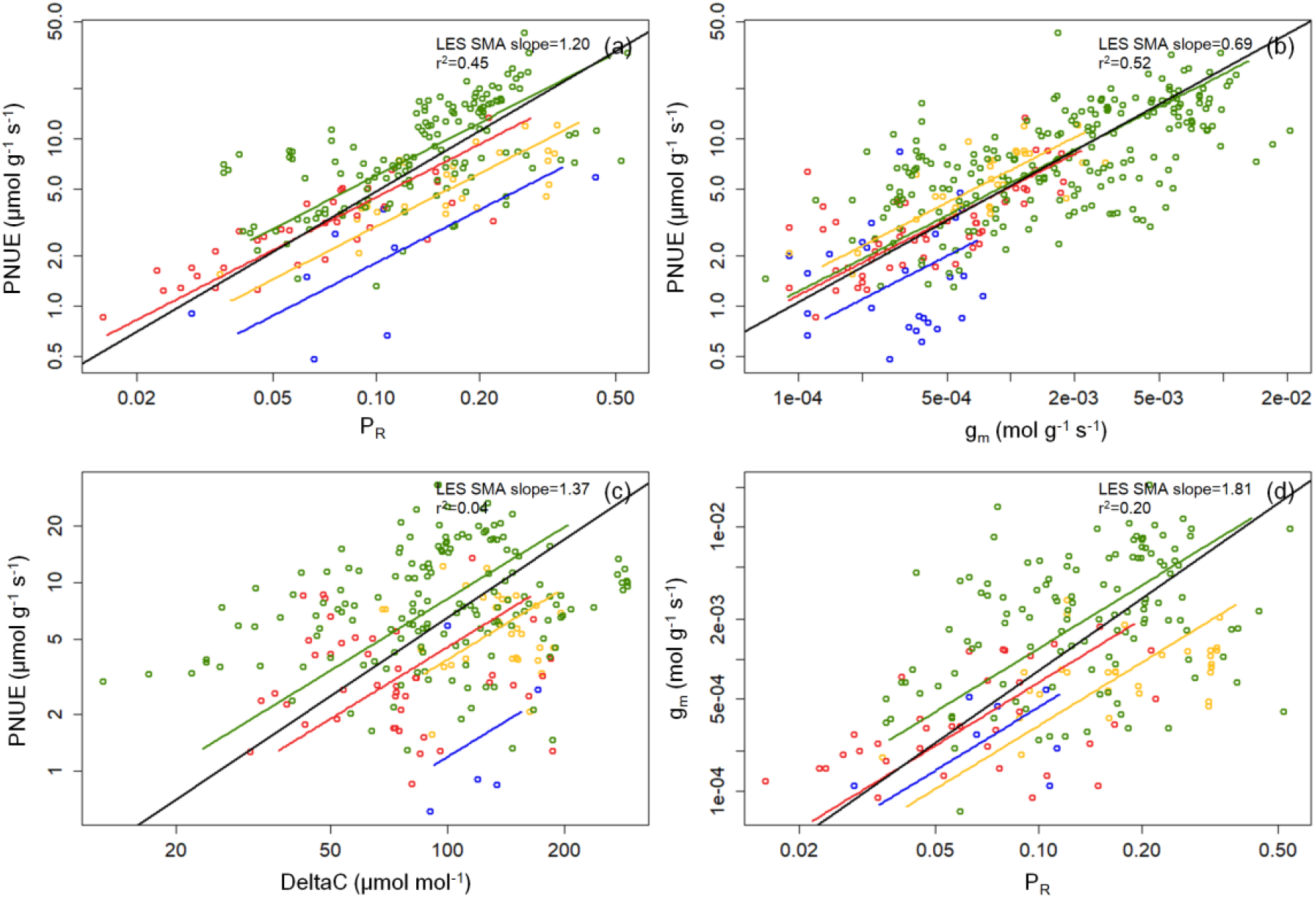
Paired-trait correlations between photosynthetic N-use efficiency (PNUE), allocation fraction of leaf nitrogen to Rubisco (P_R_), mass-based mesophyll conductance (g_m_), and internal CO_2_ drawdown (DeltaC). (a) PNUE–P_R_, (b) PNUE– g_m_, (c) PNUE–DeltaC, (d) P_R_–g_m_. The black line represents the regression of standard major analysis (SMA) fit to all data in the four phyla (LES SMA) and the other lines represent the regressions of SMA fit to all data of each phylum. The r^2^ and slope values were elucidated for the LES SMA regression.

The relative importance of P_R_ and g_m_ in determining PNUE is shown in Figure 4. The PNUE predictions corresponded to observations efficiently at both low- and high-g_m_ species assembles (Figure 4a). A close correspondence between PNUE predictions and observations was also noticed in most sampled species studied (Figure S2). The slope of the PNUE–P_R_ relationship was 57.58 in the high-g_m_ species assemble and 20.52 in the low-g_m_ species assemble. It indicated that for species with high g_m_, a 0.1-unit increase in P_R_ contributed to a 5.8-unit increase in PNUE, while the increase in PNUE was only 2.1-unit in low-g_m_ species. An exponential decreasing tendency in l_m_-l_b_ with g_m_ was observed in sampled species (Figure 4b). The l_m_ overweighed l_b_ in species with low g_m_ < 0.001 mol g^−1^ s^−1^ (approximately 0.08 mol m^−2^ s^−1^), indicating stronger mesophyll limitation on photosynthesis; conversely, species with high g_m_ > 0.001 mol g^−1^ s^−1^ exhibited a stronger biochemical limitation on photosynthesis. These results indicated a stronger limitation by g_m_ on PNUE for species with low g_m_ and a stronger limitation by P_R_ on PNUE for species with high g_m_.

**Figure 4.**
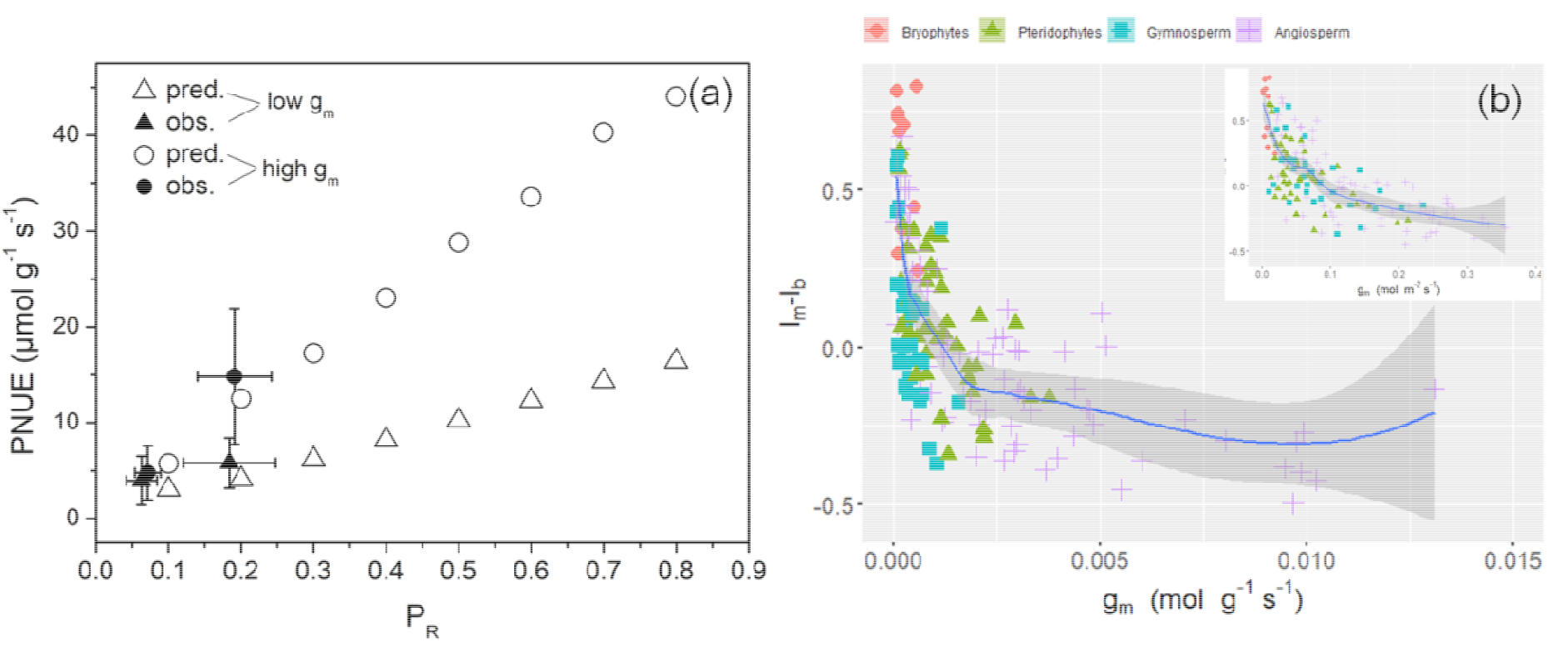
Model simulations and observations of photosynthetic N-use efficiency (PNUE) at a continuous series of Rubisco nitrogen allocation fraction (P_R_) and at two assembles of mesophyll conductance (g_m_), such as high and low ranges (circles and triangles, respectively) in subplot a. Subplot b demonstrates the relationship between l_m_-l_b_ and mass-and area-based g_m_. The high-g_m_ species assemble had a g_m_ range of 0.25–0.32 mol m^−2^ s^−1^; whereas the low-g_m_ species assemble and 0.05–0.06 mol m^−2^ s^−1^, corresponding to chloroplast CO_2_ concentration (C_c_) at approximately 100 and 230 μmol mol^−1^ under current atmospheric CO_2_ conditions, respectively. pred.: predictions; obs.: observations; l_m_-l_b_: the difference between mesophyll conductance limitation and biochemical limitation.

### Relationships between morpho-anatomical traits and P_R_ and g_m_

Relationships between morpho-anatomical traits and P_R_ and g_m_ are illustrated in Figure 5. The P_R_ exhibited a weak correlation with LMA across the four phyla (r^2^ = 0.19, Figure 5a) and the weak correlation was also observed between g_m_ and LMA (r^2^ = 0.24, Figure 5b). Statistical findings in comparison between Onoda P_R_ chemical assay dataset and the P_R_ numerical calculations in spermatophyte exhibited no significant differences in SMA slope and intercept (p = 0.1, Figure S3), indicating that the incorporation of the P_R_ numerical calculations into Onoda P_R_ chemical assay dataset did not introduce systematic biases in trait correlation analysis, and the P_R_ mixture dataset was reliable. The four phyla exhibited a consistent relationship with high correlation strength for g_m_ and T_cwm_ (r^2^ = 0.44, Figure 5c). The g_m_ in four phyla had the tendency to converge toward the Loess regression line (inset in Figure 5c). A rapid decline in g_m_ with T_cwm_ was observed, and g_m_ had the tendency to level off at higher T_cwm_. A strong correlation was observed between g_m_ and the mass-based S_c_ (r^2^ = 0.66, Figure 5d). No other anatomical traits revealed a significant correlation with g_m_ (r^2^ < 0.1, Figures S4a–f). The P_R_ was inversely proportional to T_cwm_ with an average r^2^ = 0.30 (Figure 5e). Although their correlation was much weaker (r^2^ = 0.14, Figure 5f), the DeltaC increased with increasing T_cwm_, indicating marginal effects of T_cwm_ on the internal CO_2_ drawdown.

**Figure 5.**
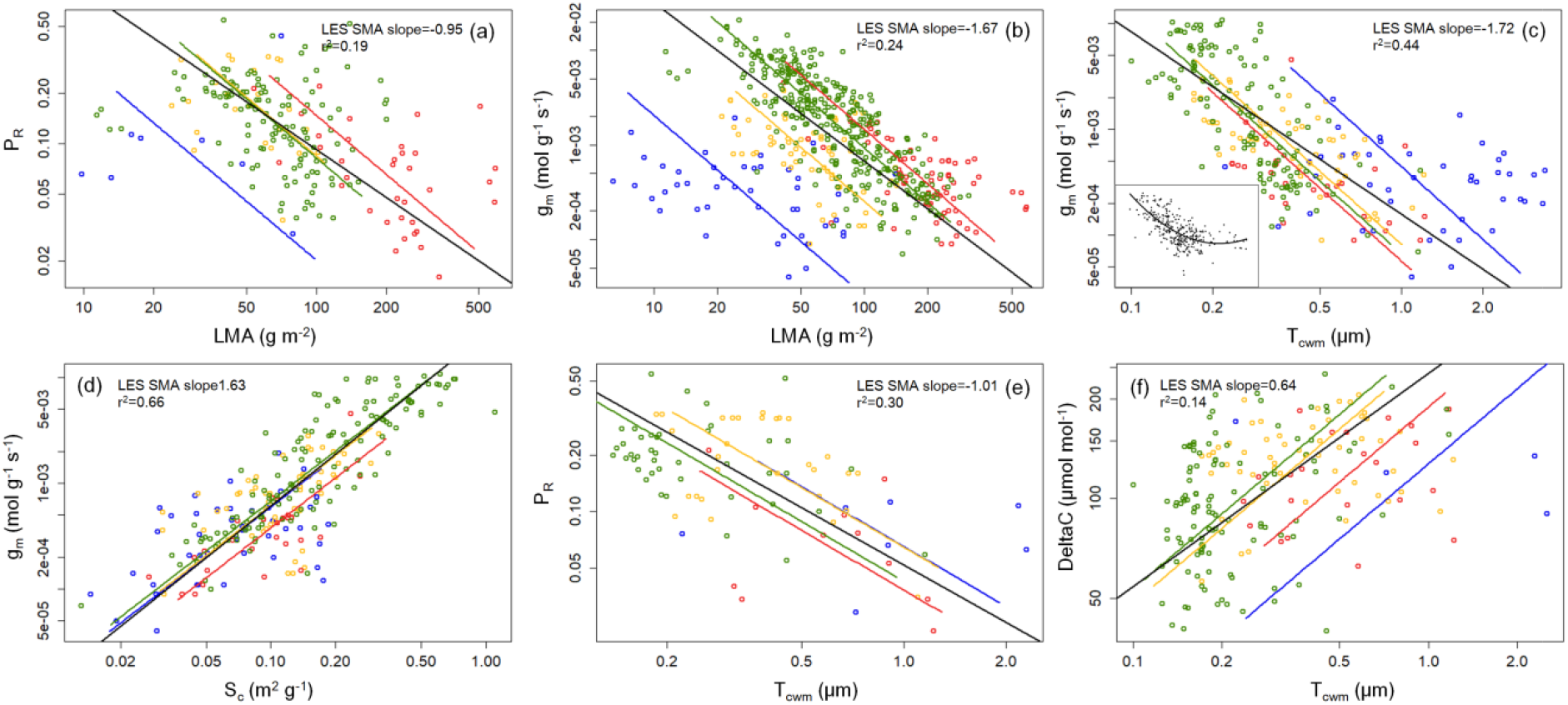
Paired-trait correlations between allocation fraction of leaf nitrogen to Rubisco (P_R_), leaf mass per area (LMA), mesophyll conductance (g_m_), mesophyll cell wall thickness (T_cwm_), chloroplast surface exposed to intercellular airspace area per unit leaf mass (S_c_), and internal CO_2_ drawdown (DeltaC). (a) P_R_–LMA, (b) g_m_–LMA, (c) g_m_–T_cwm_, (d) g_m_–S_c_, (e) P_R_–T_cwm_, (f) DeltaC–T_cwm_. The black line represents the regression of standard major analysis (SMA) fit to all data in the four phyla (LES SMA) and the other lines represent the regressions of SMA fit to all data of each phylum. The r^2^ and slope values were elucidated for the LES SMA regression. The inset in subplot c exhibited the Loess model fit to the g_m_–T_cwm_ correlation.

### Path analysis for leaf morpho-anatomical and physiological traits and PNUE

According to path analysis for the SEM fit, the seven traits, such as g_m_, P_R_, LMA, T_cwm_, mass-based S_c_, C_i_, and C_c_ together explained 88% of PNUE variations, while P_R_ and g_m_ interpreted 65% of PNUE variations (CFI = 1.0, SRMR = 0.02, p-value = 0.001, Figure 6a). The traits utilized for path analysis were chosen primarily according to the aforementioned paired-trait correlation analysis and Eqn. 3 parameters. The absolute value of path coefficient (*Q*) for T_cwm_–g_m_ was 0.84, which was higher by 13.5% than in comparison to LMA–g_m_. The S_c_–g_m_ had a high *Q* = 0.89, confirming their positive correlation shown in Figure 5d. A higher *Q* value of 27.5% was found in the T_cwm_–P_R_ in comparison to the LMA–P_R_. The *Q* values for P_R_–C_i_ or P_R_–C_c_ were less than 0.3, which suggested minimal effects of leaf internal CO_2_ concentration on P_R_ variations across species. The P_R_ and g_m_ exhibited direct and substantial effects on PNUE (*Q* > 0.45), while T_cwm_ and LMA exhibited direct and much weaker effects on PNUE (*Q* = 0.1 and 0.25, respectively). The phylogenic pattern of T_cwm_ across four phylum groups and the paired-trait correlation for PNUE– T_cwm_ are provided in Figures 6b and c. A trend of consistent decline in T_cwm_ from bryophytes to angiosperms was observed. The PNUE was negatively proportional to T_cwm_ with r^2^ as high as 0.61, which was higher by five-fold in comparison to the PNUE–LMA relationship illustrated in Figure 1c. According to the findings, P_R_ and g_m_ variations across the four phyla and through plant phylogeny were primarily related to T_cwm_. Effects of T_cwm_ on PNUE were achieved through its controls on P_R_ and g_m_.

**Figure 6.**
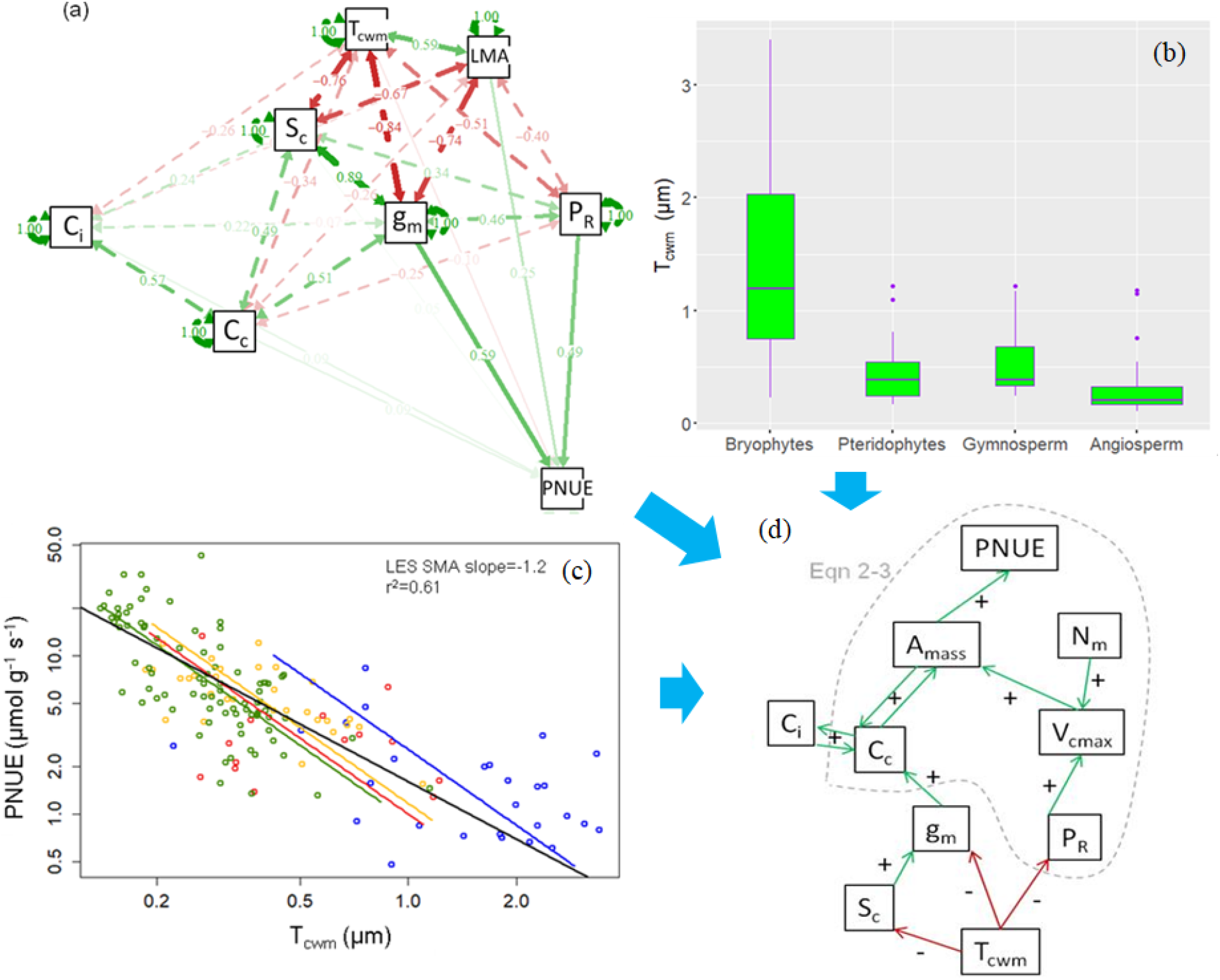
(a) The circle layout of the path diagram that indicates relationships and the causal direction between PNUE and morpho-physiological traits, (b) the phylogenetic patterns of mesophyll cell wall thickness (T_cwm_), (c) the paired-trait correlation between photosynthetic-N use efficiency (PNUE) and T_cwm_, and (d) the structural diagram about T_cwm_-mediated physiological mechanisms controlling PNUE variations. The path coefficient is represented by the arrow and value between each trait and PNUE in subplot a. The boxes were drawn with widths proportional to the square-roots of the number of observations in each phylum in subplot b. The black line represents the regression of standard major analysis (SMA) fit to all data in the four phyla (LES SMA) and the other lines represent the regressions of SMA fit to all data of each phylum in subplot c. The r^2^ and slope values were elucidated for the LES SMA regression. Interrelationships between PNUE and leaf traits embedded by the gray dashed circle in subplot d were deduced from Eqn. 2 and 3. The “+” and “-” signs reflected positive and negative relations, respectively. P_R_: allocation fraction of leaf nitrogen to Rubisco; LMA: leaf mass per area; g_m_: mass-based mesophyll conductance; T_cwm_: mesophyll cell wall thickness; S_c_: chloroplast surface area exposed to intercellular airspace per unit leaf mass; C_i_ and C_c_: leaf intercellular and chloroplastic CO_2_ concentration, respectively.

## Discussion

Thematic results with causal relationships between leaf functional traits and PNUE are illustrated in Figure 6d. According to Eqn. 2 and 3, A_mass_ is determined by C_c_ and V_cmax_, which are in turn primarily related to C_i_ and g_m_, P_R_ and N_m_, respectively. An increase in N_m_ could contribute to V_cmax_ and, subsequently A_mass_, while its contributions to PNUE were not significant (Figure S1). Indeed, A_mass_ and N_m_ were positively correlated across species (LES SMA r^2^ = 0.34). Therefore, coordinating g_m_, C_i_ and P_R_ to maximize A_mass_ per unit of leaf nitrogen is the prior biological strategy adopted by plants to obtain high PNUE in the process of evolution (Figure 6d). According to earlier research, C_i_ varied slightly among species grown under non-stress conditions in the current atmospheric CO_2_ conditions (von Caemmerer and Farquhar, 1981; Evans *et al*., 2009). In the global database, the coefficient of variation (CV) for C_i_ was 0.15, which was much lower than the CV of C_c_ (0.31). It might be reasonable to infer that g_m_, rather than g_sw_, plays a critical role in determining C_c_ across non-stressed plant species (Ethier and Livingston, 2004; Tosens *et al*., 2012; Onoda *et al*., 2017). Bryophytes have thick cell walls and no stomatal conductance structure. Therefore, it’s better to choose common leaf traits as bio-indicators to explore PNUE global variations of terrestrial plants. Cell walls (CWs) have been recognized for their importance in plant physiology and biotechnology, and recent reviews have suggested that T_cwm_ could be crucial in regulating photosynthesis (Flexas and Carriquí, 2020; Flexas *et al*., 2021). In the following sections, T_cwm_-mediated physiological mechanisms controlling PNUE variations through land plant phylogeny have been discussed.

The P_R_ and g_m_ have been considered two important factors underlying PNUE variations in seed plants (Hikosaka, 2004; Onoda *et al*., 2017). Our results demonstrated that 65% of PNUE variations in the global dataset are explained by the two variables. There are synergetic effects of the two traits on PNUE across plant phylogeny, which means an increase in either of them could contribute to the improvement of PNUE. However, the synergetic effect of P_R_ is determined by g_m_. A slight improvement in PNUE is observed in species with low g_m_ owing to an increase in P_R_; whereas a double increment in PNUE is observed in species with high g_m_ (Fig. 4a). CO_2_ enrichments inside chloroplasts owning to high g_m_ facilitate photosynthesis and subsequently a significant increment in PNUE is anticipated. Warren and Adams (2006) suggested that variation in the g_m_–photosynthesis relationship contributes to variation in PNUE. Their suggestion is supported and underlying mechanisms are addressed by our results that photosynthesis is strongly restricted by g_m_ in species with low g_m_ < 0.001 mol g^−1^ s^−1^ (Figure 4b); in contrast, the biochemical limitation is a major factor constraining photosynthesis in high-g_m_ species. Greater improvement in PNUE is achieved in high-g_m_ species than the low-g_m_ species in condition of the same P_R_. The g_m_ threshold where the relative importance of P_R_ and g_m_ in determining PNUE shifts might be 0.001 mol g^−1^ s^−1^, corresponding to the area-based g_m_ from 0.05 to 0.1 mol m^−2^ s^−1^. Xiong and Flexas (2018) evaluated the covariance in A_mass_, g_m_, and Rubisco concentrations in hundreds of rice varieties, finding low g_m_ limitation and strong biochemical limitation in modern varieties. PNUE improvement in modern varieties would be facilitated by the improvement of carboxylation capacity. A pronounced increment in PNUE would be possible in high-g_m_ species via efforts to ameliorate carboxylation efficiency. Collectively, PNUE variations across major plant groups are strongly regulated by P_R_ and g_m_, with their interactive effects varying depending on g_m_.

The g_m_ is a composite conductance of the CO_2_ diffusion pathway between the substomatal cavities and carboxylation sites within chloroplasts. Earlier studies emphasized the importance of cell wall composition (Ellsworth *et al*., 2018; Carriquí *et al*., 2020; Roig-Oliver *et al*., 2021), cell wall porosity (Evans *et al*., 2009), cell wall thickness (Carriquí *et al*., 2015, 2019, 2021; Flexas *et al*., 2021), and chloroplast distribution, in particular the S_c_ (Carriquí *et al*., 2015, 2019; Tosens *et al*., 2016; Han *et al*., 2018; Du *et al*., 2019; Ren *et al*., 2019; Nadal *et al*., 2021), in determining g_m_. Our findings indciated a strong and positive correlation between g_m_ and S_c_, while a significantly negative relationship between g_m_ and T_cwm_ across major plant groups and through phylogeny (Figures 5c and d; Figure 6a). There is an exponential decline tendency in g_m_ with increasing T_cwm_, and the low g_m_ value tends to level off at greater T_cwm_ (Figure S5), implying that species with thicker cell walls might have the same g_m_ in comparison to those with relatively thin cell walls. According to Nadal *et al*. (2021), substantial S_c_ could compensate for thick cell wall limitation to achieve relatively high g_m_ in resurrection plants with thick T_cwm_. A large range of g_m_ at given T_cwm_ in Figure 5c mainly arises from differences in S_c_ among species. The negative correlation between S_c_ and T_cwm_ across major plant groups suggests high S_c_ clustering at the low-end of T_cwm_ axis. Optimal arrangement of chloroplast position along cell walls could minimize CO_2_ diffusion resistance at liquid phase and enlarge surface area of bicarbonate ion diffusion, contributing to g_m_.

In the context of vegetation model parameterization, P_R_ global patterns were deduced from the known LMA global map using a P_R_–LMA relationship (Luo *et al*., 2021). In line with their reports, the P_R_ is inversely scaled with LMA (Figure 5a), and the relationship is analogous to the g_m_–T_cwm_ relationship. There is a progressive phylogenetic trend associated with LMA and T_cwm_ in spermatophyta: from low LMA and thin T_cwm_ in angiosperms to high LMA and large T_cwm_ in gymnosperms (Onoda *et al*., 2017; Flexas *et al*., 2021; Xing *et al*., 2021), while an exception occurred in mosses and ferns that have low LMA (Tosens *et al*., 2016; Carriquí *et al*., 2019). Interestingly, there is a consistent phylogenetic trend regarding T_cwm_ from bryophytes to angiosperms reported in the study of Gago *et al*. (2019) and in Figure 6b: from very large T_cwm_ in bryophytes to very low T_cwm_ in angiosperms. Indeed, higher correlation strength is attained when scaling g_m_ with T_cwm_ in comparison to g_m_–LMA (Figures 5b and c and Figure 6a). When g_m_ changes with T_cwm_, the log-transformed g_m_–T_cwm_ correlation exhibits more convergent characteristics (i.e., g_m_ converges at given T_cwm_). It indicated that the P_R_–T_cwm_ correlation is likely synonymous with the exponential decay correlation of P_R_–LMA, or at least ought to be. Theoretical predictions by Evans *et al*. (2009) revealed that other elements in the mesophyll CO_2_ diffusion pathway of the liquid phase could co-vary with T_cwm_, such as the decrease in Rubisco per unit of exposed chloroplast surface area and thinner chloroplast with increasing T_cwm_. If the P_R_, which is proportional to photosynthesis, did not decrease with increases in T_cwm_, then the internal CO_2_ drawdown would have been substantially higher than observations. Assuming independence of P_R_ on T_cwm_, the ten-fold decline of g_m_ owning to an increase in T_cwm_ from 0.2 to 0.5 μm (e.g., the g_m_ decline from 0.004 to 0.0004 mol g^−1^ s^−1^) would amplify the internal CO_2_ drawdown by more than five folds in species featured by T_cwm_ of 0.5 μm, and a strong correlation between DeltaC and T_cwm_ would therefore be observed. However, this expectation contradicts the fact that the actual increase of the CO_2_ drawdown is limited to 50% over a wide range of T_cwm_ (Figure 5f). The decrease in P_R_ with T_cwm_ ultimately balances the increase in mesophyll resistance, resulting in the CO_2_ drawdown that is only weakly proportional to T_cwm_ (Figure 5f). The experimental results in this study, i.e., a significantly negative correlation between T_cwm_ and P_R,_ corroborate the theoretical prediction of the T_cwm_–biochemical capacity offset mechanisms in the study of Evans *et al*. (2009) (Figure 5e and Figure 6a). In our study, the observed significance of the P_R_–T_cwm_ relationship across major plant groups is consistent with the findings in the study by Onoda *et al*. (2017) for seed plants. However, the anticipated inverse correlation between chloroplast thickness and T_cwm_ is not observed (Figure S6). The thickness of photosynthetic tissues is one of the distinctive differences in leaf anatomy among bryophytes, ferns, and spermatophytes (Han *et al*., 2018; Carriquí *et al*., 2019). The chloroplast amount per leaf mass may co-vary with T_cwm_ from bryophytes to angiosperms, resulting in the observed P_R_–T_cwm_ relationship. It was unclear and open to interpretation whether the response pattern of P_R_ to T_cwm_ was exclusively independent or dependent causality owing to g_m_ changes. The possibility of g_m_ effects on the observed P_R_–T_cwm_ response pattern could be excluded as no strong correlations are observed for the P_R_–g_m_ relationship (Figure 3d) and the V_cmax_–C_c_ relationship (Figure S7). It was proposed that T_cwm_ is the critical factor controlling evolutionary modifications of PNUE from bryophytes to angiosperms based on the generality of significant correlations for P_R_–T_cwm_ and g_m_–T_cwm_ in the worldwide database.

Mosses and ferns are widely distributed across the globe, being dominant species in various habitats ranging from understory to full sunlight environment and from hot and moist lands to cold and dry lands. A recent study demonstrated that the paired-trait correlations in leaf economics spectrum in ferns are independent of growth light and nutrient gradients (Li *et al*., 2022). Results in 75 woody species in subtropical forests in the study by Chen *et al*. (2020) revealed a unified trade-off relationship in paired traits under different light gradients. Differences in between-species responses to changes in light conditions in trees and shrubs arise from significant shifts in the intercept of the SMA regression line not the scaling slope (Keenan and Niinemets, 2016), which might be explained by either leaf habits or the height-based economics spectrum (Zhao *et al*., 2017). Species-level acclimations to habitat irradiance could not be strong enough to change the relative positions of moss species along trait axes (Waite and Sack, 2009; Z. Wang *et al*., 2017). Most deviations of bryophytes in A_mass_–LMA and PNUE–LMA relationships in comparison to angiosperms are attributed to fundamental difference between the two plant groups. Our results concerning underpinnings of PNUE variations from bryophytes to angiosperms are reliable.

## Materials and Methods

### Data collection

In this study, we constructed a global collection of leaf morphological, anatomical, and physiological traits of non-stressed mature leaves of terrestrial C_3_ species comprising major plant groups and phylogeny. The dataset included the photosynthetic organs with various morphologies (e.g. phyllidia or thalli in mosses and lichen species). LMA and leaf thickness (T_leaf_), chloroplast surface area exposed to intercellular airspace per unit leaf area (S_c_), mesophyll surface area exposed to intercellular airspace per unit leaf area (S_m_), cytoplasm thickness (T_cyt_), mesophyll cell wall thickness (T_cwm_), mesophyll thickness (T_mes_), chloroplast thickness (T_chl_), the fraction of intercellular air space (ƒ_ias_), the average distance between the chloroplast and the cell wall (cytoplasm thickness, L_ct_) were among the structural and anatomical traits of the leaf. Physiological traits comprised photosynthetic capacity at normal atmospheric CO_2_ concentration and saturation light (A_mass_, mass-based photosynthesis rate), leaf nitrogen content per leaf mass (N_mass_), stomatal conductance to water vapor (g_sw_), intercellular and chloroplastic CO_2_ concentration (C_i_ and C_c_, respectively), g_m_, the C_c_-based maximum carboxylation rate (V_cmax_), the maximum electron transport rate (J_max_), stomatal limitation on photosynthesis (l_s_), mesophyll limitation on photosynthesis (l_m_), biochemical limitation on photosynthesis (l_b_), and leaf temperature (L_temp_). In this study, articles that highlighted keywords, such as “mesophyll conductance” or “internal conductance” archived by the Web of Science (https://www.webofscience.com) and the China National Knowledge Infrastructure (https://www.cnki.net) until July 2022 were considered. Trait measurements in most species including ferns and moss species in the field and/or outdoor conditions under the most favorable growing conditions (i.e., with the least amount of environmental stress), were considered reliable for use. Measurements of shade-adapted leaves within plant canopies were not included in the global dataset.

The data collection campaign was challenging as the number of available traits varied among surveyed articles. If the traits could be inferred by supporting information in articles, the inferred values were restored to the global database. Indeed, it is impossible to determine the CO_2_ concentration within the chloroplast stroma. The internal CO_2_ drawdown (DeltaC, the difference between C_i_ and C_c_) was inferred according to Fick’s first law: DeltaC = C_a_ – A(1.56/g_sw_ – 1/g_m_), which was commonly adopted in leaf photosynthesis-stomatal conductance coupled model (Onoda *et al*., 2017; Xue *et al*., 2022). It was discovered that bryophytes are the only group of plants that are completely devoid of stomata. Hence, the DeltaC values in bryophytes species were inferred using the equation of, DeltaC = A/g_m_. The global C_a_ of 380 ppm was applied in the absence of C_a_ data in surveyed articles.

The P_R_ was calculated using two different methods: a numerical calculation method using V_cmax_ and N_area_ as demonstrated by Niinemets and Tenhunen (1997), and a chemical assay of Rubisco content. The Sharkey online calculator (Sharkey *et al*., 2007) and optimization inversion algorithms were two common methods to estimate V_cmax_ from the A/C_i_ curve (Xue *et al*., 2022). The estimated V_cmax_ values by the parameter estimation methods were comparable among plant species, since Rubisco kinetics parameters in 89% of surveyed articles were consistently taken from the study of Bernacchi *et al*. (2002). Variation in Rubisco kinetics parameters measured *in vitro* was less than 10% among C_3_ plant species (von Caemmerer, 2020), which was far less than the 10-fold variations of inter-specific A_mass_ at a given LMA (Figure 1d). Hence, their Rubisco kinetics parameters might be highly recommended for V_cmax_ calculations across major plant functional types, as commonly did in surveyed articles. The P_R_ dataset derived from the chemical assay in the study of Onoda *et al*. (2017) was solely limited to spermatophyte. By combing P_R_ numerical calculations, the P_R_ dataset was expanded to enrich species diversity in the global dataset. In the global dataset, all leaf traits were measured in sunlit leaves around 25°C with relatively high incident radiation and ambient atmospheric CO_2_ concentration. The P_R_ assessments by the numerical method were sensitive to measuring temperatures (Niinemets and Tenhunen, 1997). According to Medlyn *et al*. (2002), temperature response functions of V_cmax_ and J_max_ with activation energy (Δ*H*_a_) at 63.0 kJ mol^−1^ were used to convert the values of V_cmax_ and J_max_ measured at other leaf temperatures to values at standard temperature.

Except for the DeltaC and P_R_, other values of physiological and morpho-anatomical traits were directly retrieved from the surveyed articles. To prevent the impact of environmental stress on analysis, leaf trait data from the same species reported by various studies were averaged for use when environmental conditions sensed by the species were optimal (Poorter *et al*., 2009; Poorter *et al*., 2011; Xing *et al*., 2021). Wild, old and modern crop kinds differed significantly in leaf morpho-physiological traits, being representatives of angiosperm species in leaf economics spectrum evolving towards high photosynthetic efficiency (Xiong and Flexas, 2018; Martin *et al*., 2018; Hayes *et al*., 2019). In such case, physiological and morpho-anatomical traits in crops and non-crop plants were merged for data analysis. Collectively, there were 1296 lines of data in 679 species, covering major life forms in land plants, such as bryophytes (57 lines in mosses and lichen species), pteridophytes (64 lines in ferns and fern allies species), gymnosperm (83 lines in needles and scales leaved species), and angiosperm (1092 lines in trees, shrubs, grass and crops).

### Data analysis

Data of all traits were log_10_-transformed before correlation analysis, which was a common practice adopted in ecological studies because data types of some leaf traits do not follow a normal distribution (Wright *et al*., 2005; Carriquí *et al*., 2015; Tosens *et al*., 2016; Onoda *et al*., 2017; Carriquí *et al*., 2019). The strength of bivariate trait correlation was assessed using the standardized major axis slope (SMA, also known as reduced major axis slopes), a method frequently practiced in these studies, was applied to determine. The non-parametric regression model (locally weighted regression, Loess) used by Xue *et al*. (2021) and Kurmangozhinov *et al*. (2020) to clarify threshold at which trait strong coupling shifted to weak coupling was used for paired traits with significant curvilinear relationship, rise to the maximum or decline to the minimum curve trajectory. Indeed, it is the mass-based physiological traits, such as A_mass_ that determine the biochemical capacity of single cells and are the important factors in the global trade-off between the physiological and structural traits of foliage (Niinemets and Tenhunen, 1997; Wright *et al*., 2005; Tosens *et al*., 2016; Onoda *et al*., 2017). We found that the area-based trait analysis exhibited analogous results in correlation analysis as the expression of the mass-based traits. Thus, the mass-based physiological traits were employed for correlation analysis unless otherwise indicated. Prior to paired trait correlation analysis, a phylogenetic tree was constructed for all species using the megatree approach in the R package V.PhyloMaker and phylogenetic signals were estimated using Pagel’s λ following Xing *et al*. (2021). There were 39 species failing to be bound to the constructed phylogenetic tree as they were not present in the plantlist database of the V.PhyloMaker package (package acquiring date: 2022/05/13). Of the 39 species, *Polistichium aculeatum* (pteridophyte) was one, and the remaining belonged to the bryophytes.

In R 4.0.5 software (R Core Team, 2021), the paired-trait correlation analysis using the smart-package, Loess model application, and phylogenetic signal analysis using the V.PhyloMaker package were performed. Structural equation modeling (SEM) was applied to explore how morpho-physiological traits influence PNUE variations (Shipley, 2004). The estimator to parameterize the SEM was the maximum likelihood method. Using the comparative fit index (CFI, > 0.9), p-value, and the standard root mean squared residual (SRMR, < 0.08) (Shipley, 2004), appropriate SEM model fit was inspected. The semPlot package in the R software was used to visualize the SEM analysis that was performed using the lavaan package (Rosseel, 2012).

The relative importance of P_R_ and g_m_ in relation to PNUE variations was quantified using the photosynthetic coordination theory. According to the biochemical photosynthesis model established by Farquhar *et al*. (1980), the photosynthetic rate is constrained by either the Rubisco carboxylation rate at low Rubisco content and CO_2_ concentrations (A_c_) or the RuBP regeneration rate at comparatively high CO_2_ concentrations owning to low electron transport rates (A_j_). The A_c_ value is primarily determined by V_cmax_, followed by C_c_ at a given leaf temperature, (Eqn. 1). Besides the direct effects of P_R_ on photosynthesis, the effects of g_m_ on photosynthesis are realized through their impacts on C_c_ (Ethier and Livingston, 2004; Evans *et al*., 2009; Kaldenhoff, 2012; Tosens *et al*., 2012; Onoda *et al*., 2017; Gómez-Ocampo *et al*., 2021). The photosynthetic coordination theory was originally proposed by von Caemmerer and Farquhar (1981) and further modified by Maire *et al*. (2012) in 31 individual C_3_ species. H. Wang *et al*. (2017), Smith *et al*. (2019), and Harrison *et al*. (2021) in regional vegetation studies stated that photosynthesis tends to be co-limited by electron transport and carboxylation under typical environmental conditions, thus maximizing potential resource use for growth and reproduction. The P_R_ and g_m_ functioning effects on inter-specific PNUE variations could be predicted quantitatively using the photosynthetic coordination theory. The PNUE could be expressed with Eqn. 3 that was derived from Eqn. 1 and 2 as follows:

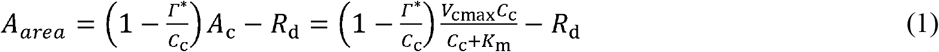

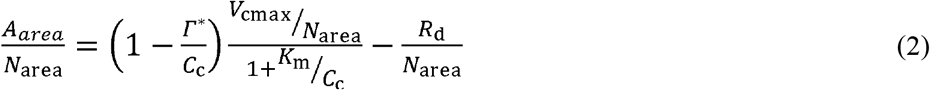

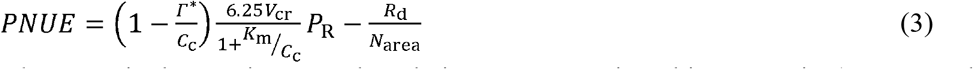

where V_cr_ is the maximum carboxylation rate per unit Rubisco protein (47.34 μmol CO_2_ (g Rubisco)^−1^ s^−1^ at 25°C) and 6.25 is the nitrogen content of Rubisco protein (Niinemets and Tenhunen, 1997; Luo *et al*., 2021); R_d_ is area-based dark respiration in the light; Γ* is the chloroplast CO_2_ compensation point, corresponding to 42.75 μmol mol^−1^ at 25°C. K_m_ is equal to K_c_(1+O/K_o_), where K_c_ and K_o_ are the Michaelis– Menten constants for CO_2_ and O_2_, respectively (Bernacchi *et al*., 2002). The PNUE simulations were generated considering a continuous series of P_R_ and C_c_. We hypothesized that the effect of g_m_ on PNUE were more obvious across contrasting plant groups with significant differences in g_m_ in comparison to several randomly selected species. As a result, the species in the global dataset were divided into two groups according to their g_m_ values, such as the high- and low-g_m_ species assembles. The high-g_m_ species assemble exhibited g_m_ from 0.25 to 0.32 mol m^−2^ s^−1^, and the low-g_m_ species assemble had g_m_ from 0.05 to 0.06 mol m^−2^ s^−1^, corresponding to C_c_ at approximately 100 and 230 μmol mol^−1^ in leaves under current atmospheric CO_2_ conditions, respectively. The PNUE theoretical analysis was cross-validated using the PNUE and P_R_ measurements together with g_m_ and C_c_ measurements (i.e., trait values taken/inferred from the surveyed articles) in the two species assembles. Variations in predicted PNUE were assumed to be independent of the intercept R_d_/N_area_.

## Acknowledgments

This work was supported by the National Natural Science Foundation of China (32001129) and Estonian Research Council (PUT 1473).

## Author contributions

Conceptualization and investigation: DDL, WX

Methodology, supervision, visualization and Writing-original draft: WX

Writing-review & editing: MC, DLX, TT, YCX, WX

## Data and materials availability

All data are available in the supplementary materials.

